# Effect of C-to-T transition at CpG sites on tumor suppressor genes in tumor development in cattle evaluated by somatic mutation analysis in enzootic bovine leukosis

**DOI:** 10.1101/2024.03.19.585713

**Authors:** Asami Nishimori, Kiyohiko Andoh, Yuichi Matsuura, Tomohiro Okagawa, Satoru Konnai

## Abstract

Oncogenic transformation of normal cells is caused by mutations and chromosomal abnormalities in cancer-related genes. Enzootic bovine leukosis (EBL) is a malignant B-cell lymphoma caused by bovine leukemia virus (BLV) infection in cattle. Although a small fraction of BLV-infected cattle develops EBL after a long latent period, the mechanisms for oncogenesis in EBL cattle remain largely unknown. In this study, we analyzed the types and patterns of somatic mutations in cancer cells from 36 EBL cases, targeting 21 cancer-related genes. Various somatic mutations were identified in 8 genes, *TP53*, *NOTCH1*, *KMT2D*, *CREBBP*, *KRAS*, *PTEN*, *CARD11*, and *MYD88*. In addition, *TP53* gene was found to be mutated in 69.4% of EBL cases, with most being biallelic mutations. In some cases, associations were observed between the ages at which cattle had developed EBL and somatic mutation patterns; young onset of EBL possibly occurs due to congenital mutations, high impact mutations affecting protein translation, and biallelic mutations. Furthermore, nucleotide substitution patterns indicated that cytosine at CpG sites tended to be converted to thymine in many EBL cases, which was considered to be the result of spontaneous deamination of 5-methylctosine. These results demonstrate how somatic mutations have occurred in cancer cells leading to EBL development, thereby explaining its pathogenic mechanism. These findings will contribute to a better understanding and future elucidation of disease progression in BLV infection.

**Importance:** Enzootic bovine leukosis (EBL) is a malignant and lethal disease in cattle. Currently, there are no effective vaccines or therapeutic methods against bovine leukemia virus (BLV) infection, resulting in severe economic losses in livestock industry. This study provides a renewed hypothesis to explain the general mechanisms of EBL onset by combining the previous finding that several integration sites of BLV provirus can affect the increase in survival and proliferation of infected cells. We demonstrate that two additional random events are necessary for oncogenic transformation in infected cell clones, elucidating the reason why only few infected cattle develop EBL. Further exploration of somatic mutation and BLV integration sites could support this hypothesis more firmly, potentially contributing to the development of novel control methods for EBL onset.

## Introduction

Cancer is a disease of cell proliferation caused by chromosomal aberrations and somatic mutations in genes related to proliferation, the cell cycle, and DNA damage repair. Currently, at least 100 oncogenes and at least 30 tumor suppressor genes have been identified [1]. Oncogenes are considered to enhance cell proliferation by gain-of-function mutations, whereas tumor suppressor genes such as *RB1* and *TP53* cause loss of control over normal cell growth and differentiation by loss-of-function mutations or deletion [1, 2]. Generally, both copies of tumor suppressor genes on homologous chromosomes need to be inactivated to achieve oncogenic transformation, which is a concept known as Knudson’s “two-hit” theory [3]. The first “hit” is not necessarily a somatic mutation; Li-Fraumeni syndrome is a hereditary and familial cancer caused by a germline mutation in *TP53* gene. Patients with Li-Fraumeni syndrome have only one active *TP53* gene, leading to the development of multiple types of tumors at a young age due to a somatic mutation as the second “hit” in the normal allele [4, 5]. Thus, analyzing somatic mutations in cancer cells is important for understanding the history and process of oncogenesis.

Enzootic bovine leukosis (EBL) is a malignant B-cell lymphoma in cattle caused by bovine leukemia virus (BLV) infection, which belongs to the family *Retroviridae*, genus *Deltaretrovirus* [6]. BLV is closely related to human T-cell leukemia virus type I (HTLV-1) that infects human T cells and causes adult T-cell leukemia/lymphoma (ATL). BLV and HTLV-1 exhibit similar life cycles and disease progression. Their viral genomes are integrated into the host genome as a provirus, and only a small fraction (1–5%) of infected individuals develops lymphoma after a long latent period [6]. Experimental infection with BLV in cattle has shown two types of viral propagation during the infection *in vivo*. BLV actively replicates and propagates by infecting new lymphocytes in the early phase of infection (infectious cycle), but it expresses few viral antigens after the development of host acquired immunity and propagates via clonal expansion of infected cells in the chronic stage (mitotic cycle) [7]. Because infecting new target cells results in generating a new clone of infected cells with a different provirus integration site, the infectious cycle is important to increase the clonal variation of infected cells *in vivo*. A recent study reported that clonal variation and provirus integration sites were involved in the mechanisms of EBL and ATL development [8]. Rosewick et al. reported that BLV/HTLV-1 proviruses in tumor genomes of EBL animals or ATL patients were preferentially integrated near cancer driver genes, and that the proviruses could disturb mRNA expression of host genes located in their vicinity by gene interruption, depending on viral 5’ long terminal repeat poly-(A) signal or expression of 3’ antisense RNA-dependent chimeric transcript. However, the provirus integration sites did not fully explain the reasons why only a small fraction of infected individuals developed EBL or ATL, because the same pattern of integration sites was observed in animals at a polyclonal non-malignant stage [8]. The authors discussed in their report that provirus integration sites would contribute to initial selection of viral-infected cell clones at an early aleukemic stage, and the accumulation of further somatic mutation in the host genome was needed for the onset of lymphoma. Therefore, mutation analysis of cancer-related genes will enable the more detailed mechanisms of EBL development to be clarified.

To date, many studies of the mutation analyses of bovine *TP53* gene have been reported using clinical samples or cell lines derived from EBL cattle [9–13]. However, unlike the research on ATL [14], there is no information on the mutation status of other cancer-related genes involved in EBL development. Moreover, although the analysis of mutation signatures, which is an *in silico* method to evaluate the mutation process of oncogenesis from the pattern of single nucleotide variants (SNVs), is used in the field of human cancer research [15, 16], neither mutational patterns nor evaluation of allelic variation based on the “two-hit” theory has been fully elucidated in cancer cells of EBL cattle. What is more, recently, EBL onset in young cattle (less than 3 years old) has been frequently reported in Japan, despite the common ages of EBL development for BLV-infected cattle being approximately 5–8 years [17, 18]. Examining the relation between cattle ages when EBL developed and patterns of somatic mutations in cancer cells will generate much interest. Therefore, in this study, a somatic mutation analysis targeting the selected 21 genes was performed to clarify genomic changes related to oncogenesis in EBL cattle. The pattern of identified somatic mutations was then evaluated based on the ages of EBL development, and the mutation process of oncogenesis was evaluated using mutation signature analysis.

## Results

### Cancer-related genes in which somatic mutations were detected in EBL cases

According to the procedures shown in Fig. 1, somatic mutation analyses were conducted for 36 EBL cases. The results showed that somatic mutations were detected in coding sequence (CDS) regions of only 8 of 21 targeted cancer-related genes, *TP53*, *NOTCH1*, *KMT2D*, *CREBBP*, *KRAS*, *PTEN*, *CARD11*, and *MYD88* (Tables 1, 2 and S3; summarized in Fig. 2A). There were identical mutations in different EBL cattle in *TP53* (p.Arg167His, p.Arg241Trp, p.Arg242Trp p.Arg242Gln, and p.Arg275His) and *NOTCH1* (p.Cys468fs) genes, implying the presence of mutational hotspots in these genes (Table 2). *TP53* was the most frequently mutated gene, with 69.4% (25/36) of EBL cases possessing at least one *TP53* mutation in the cancer cells (Fig. 2B). Moreover, 88.0% (22/25) of *TP53* mutations were detected as biallelic mutations, namely multiple mutations or mutations associated with loss of heterozygosity (LOH) (Fig. 2A). Although the mutation frequencies of *NOTCH1* and *CREBBP* genes were not high, they were the only two genes in which biallelic mutations were observed in EBL cases with wildtype *TP53* gene, such as cases #5, #29, and #34 (Fig. 2A,B). It should be noted that *TP53* mutations with LOH in several cases were validated by comparisons between variant allele frequencies (VAFs) of blood and tumor samples for germline mutations located on chromosome 19 (Table S4).

**Figure 1.**
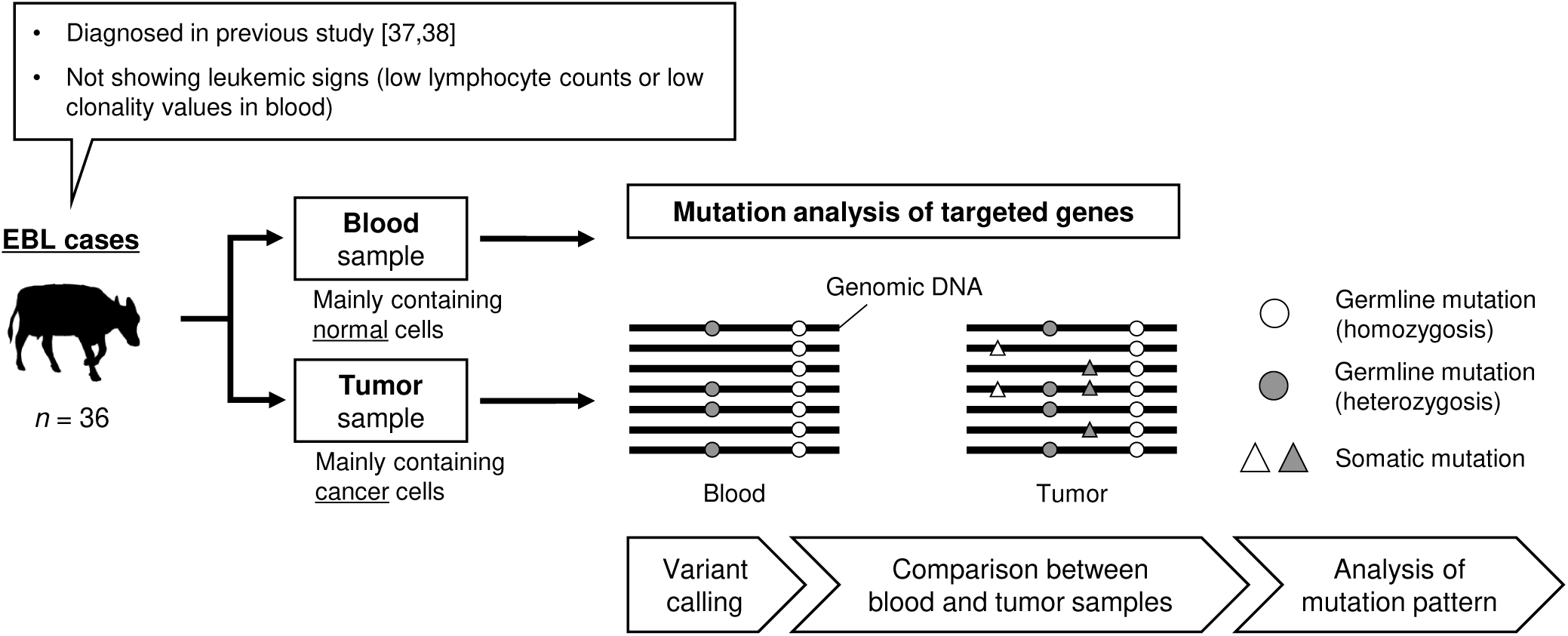
A scheme for somatic mutation analysis in cattle with EBL development. Thirty-six EBL cattle collected in previous studies [37, 38] were used for the mutation analysis. Blood and tumor samples derived from the same EBL cases were examined to identify variants in their genome using amplicon sequencing that targeted 21 cancer-related genes. Following variant calling, variant lists were compared between blood and tumor samples, and the variants existing only in tumor samples were identified as somatic mutations. All identified somatic mutation were used for mutation pattern analysis.

**Figure 2.**
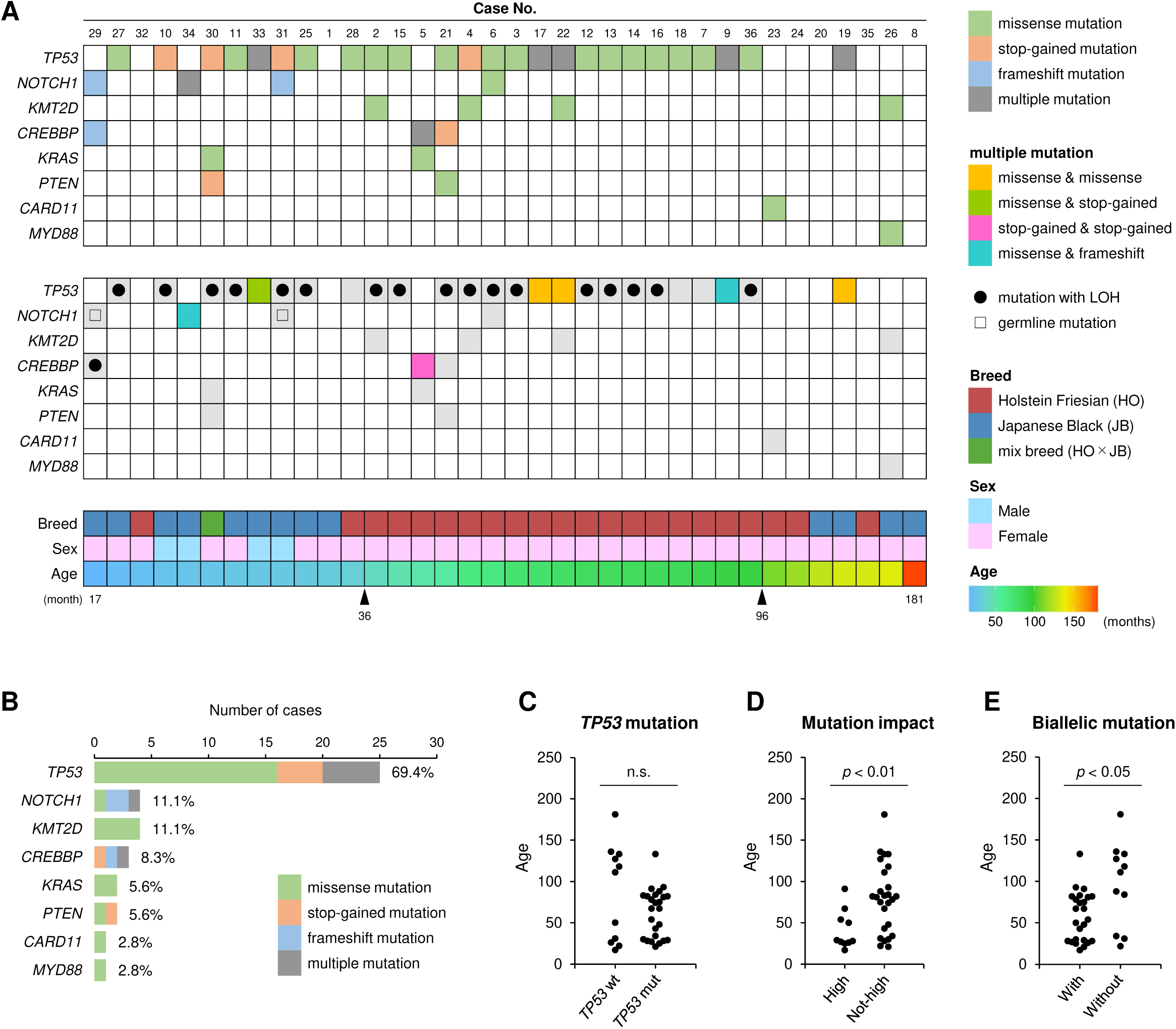
Somatic mutation analysis in 36 EBL cases at various ages in months. (A) Identified somatic mutations of 8 of 21 targeted cancer-related genes are shown in ascending order of age for EBL cases. Genes in which a somatic mutation are not identified at all are omitted. The top panel indicates mutation type, and the middle panel indicates detailed classification of multiple mutations and presence or absence of loss of heterozygosity (LOH) and germline mutations. The bottom panel indicates breed, sex, and age in months of each case. Information on identified somatic mutations based on each targeted gene or each EBL case is listed in Table 2 or Table S3, respectively. (B) Number of cases and frequencies and types of somatic mutations are shown. (C–E) Ages of cattle are compared among 36 EBL cases based on (C) presence of *TP53* mutation, (D) intensities of mutation impact, and (E) presence of biallelic mutation. *TP53* wt, wildtype *TP53*; *TP53* mut, mutated *TP53*. High mutation impact means stop-gained or frameshift mutation. Biallelic mutation includes multiple mutations or mutation with LOH. Significant differences between the two groups were evaluated by Mann-Whitney’s U test (*P*-value < 0.05).

**Table 1.**
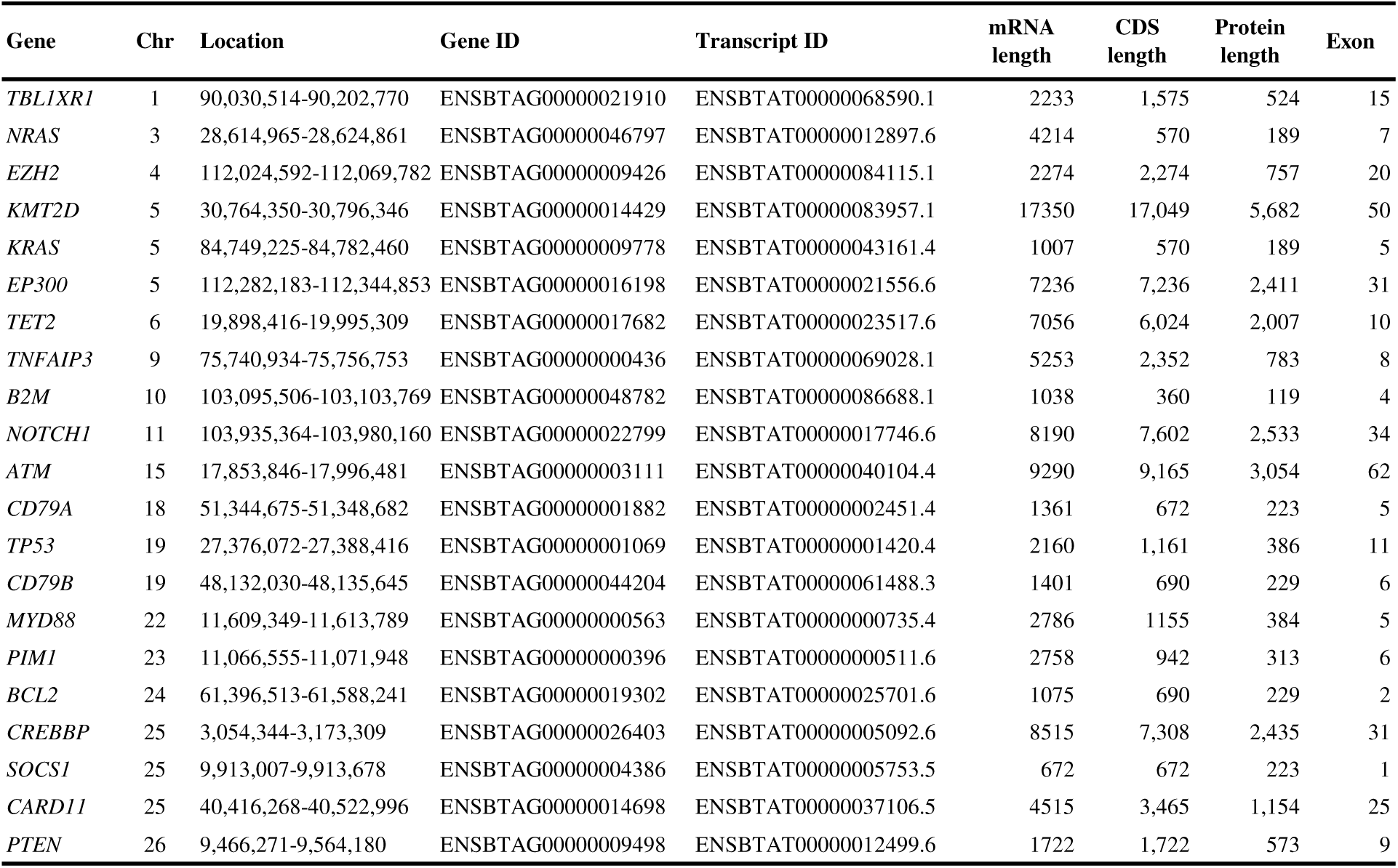
Targeted cancer-related genes in the bovine genome.

**Table 2.** Somatic mutations of 21 cancer-related genes identified from 36 EBL cases.

| Gene | Chr | Mutation type | Genome location | Exon | CDS | Amino acid substitution | Mutation contexts | No. of EBL cases | Previous reports |
| --- | --- | --- | --- | --- | --- | --- | --- | --- | --- |
| *TP53* | 19 | stop gained | 27380089 | 4/11 | c.175G>T | p.Glu59* | T[C>A]A | 33 | - |
|  |  | stop gained | 27380015 | 4/11 | c.249G>A | p.Trp83* | G[C>T]C | 4 | - |
|  |  | missense | 27379974 | 4/11 | c.290G>A | p.Gly97Asp | G[C>T]C | 36 | - |
|  |  | stop gained | 27379449 | 5/11 | c.382C>T | p.Gln128* | C[C>T]A | 30 | - |
|  |  | missense | 27379400 | 5/11 | c.431C>T | p.Pro144Leu | C[C>T]G | 33 | [9] |
|  |  | missense | 27379338 | 5/11 | c.493G>C | p.Val165Leu | A[C>G]A | 17 | - |
|  |  | missense | 27379331 | 5/11 | c.500G>A | p.Arg167His | A[C>T]G | 2, 7, 13, 16, 19 | [9–11] |
|  |  | missense | 27379319 | 5/11 | c.512A>G | p.His171Arg | A[T>C]G | 19 | - |
|  |  | missense | 27379193 | 6/11 | c.558C>G | p.His186Gln | A[C>G]C | 22 | - |
|  |  | missense | 27378842 | 7/11 | c.687C>A | p.Phe229Leu | T[C>A]A | 14 | [9] |
|  |  | missense | 27378819 | 7/11 | c.710G>A | p.Gly237Glu | C[C>T]C | 6 | - |
|  |  | missense | 27378808 | 7/11 | c.721C>T | p.Arg241Trp | C[C>T]G | 12, 18 | [9, 11, 12] |
|  |  | missense | 27378805 | 7/11 | c.724C>T | p.Arg242Trp | G[C>T]G | 11, 21 | [9, 12] |
|  |  | missense | 27378804 | 7/11 | c.725G>A | p.Arg242Gln | C[C>T]G | 9, 15 | [9, 10] |
|  |  | missense | 27378780 | 7/11 | c.749T>C | p.Leu250Pro | C[T>C]G | 28 | - |
|  |  | stop gained | 27378778 | 7/11 | c.751G>T | p.Glu251* | T[C>A]C | 10 | - |
|  |  | frameshift | 27378769 | 7/11 | c.758_759delCT | p.Ser253fs | - | 9 | - |
|  |  | missense | 27378376 | 8/11 | c.824G>A | p.Arg275His | G[C>T]G | 3, 22, 25 | [9] |
|  |  | missense | 27378365 | 8/11 | c.835G>A | p.Glu279Lys | T[C>T]C | 27 | - |
|  |  | missense | 27377694 | 10/11 | c.988C>T | p.Arg330Cys | A[C>T]G | 17 | [9] |
|  |  | stop gained | 27377679 | 10/11 | c.1003C>T | p.Arg335* | C[C>T]G | 31 | - |
| *NOTCH1* | 11 | frameshift | 103956330 | 8/34 | c.1403delG | p.Cys468fs | - | 29<sup>†</sup>, 31<sup>†</sup>, 34 | - |
|  |  | missense | 103956322 | 8/34 | c.1412A>G | p.Gln471Arg | C[T>C]G | 34 | - |
|  |  | missense | 103946432 | 20/34 | c.3248G>A | p.Arg1083His | G[C>T]G | 6 | - |
| *KMT2D* | 5 | missense | 30777496 | 25/50 | c.6344G>A | p.Arg2115Gln | C[C>T]G | 22 | - |
|  |  | missense | 30792244 | 45/50 | c.16073G>T | p.Gly5358Val | T[C>A]C | 4 | - |
|  |  | missense | 30793539 | 46/50 | c.16408G>A | p.Val5470Met | A[C>T]G | 2 | - |
|  |  | missense | 30796297 | 50/50 | c.17000C>T | p.Ser5667Phe | T[C>T]C | 26 | - |
| *CREBBP* | 25 | stop gained | 3147752 | 2/31 | c.733C>T | p.Gln245* | C[C>T]A | 5 | - |
|  |  | frameshift | 3085919 | 14/31 | c.2650dupC | p.Gln884fs | - | 29 | - |
|  |  | stop gained | 3078462 | 17/31 | c.3283C>T | p.Gln1095* | C[C>T]A | 21 | - |
|  |  | stop gained | 3061108 | 27/31 | c.4480C>T | p.Gln1494* | T[C>T]A | 5 | - |
| *KRAS* | 5 | missense | 84753447 | 2/5 | c.34G>A | p.Gly12Ser | C[C>T]A | 5 | - |
|  |  | missense | 84753447 | 2/5 | c.34G>T | p.Gly12Cys | C[C>T]A | 30 | - |
| *PTEN* | 26 | missense | 9466850 | 1/9 | c.580G>A | p.Asp194Asn | T[C>T]T | 21 | - |
|  |  | stop gained | 9538357 | 5/9 | c.898C>T | p.Arg300* | A[C>T]G | 30 | - |
| *CARD11* | 11 | missense | 40519575 | 22/25 | c.2845G>A | p.Asp949Asn | T[C>T]C | 23 | - |
| *MYD88* | 22 | missense | 11611445 | 3/5 | c.881C>G | p.Ser294Cys | T[C>G]C | 26 | - |

The symbol (†) indicates that the variant was called a germline mutation.

### Age-associated characteristics of somatic mutations

One remarkable point in the associations between the ages at which cattle had developed EBL and somatic mutation patterns is evident in cases #29 and #31, which involved young cattle less than 36 months old (Table S1). Both normal and cancer cells in these cases possessed a frameshift deletion in *NOTCH1* gene that was not registered as a known Variant ID (Fig. 2A and Table 2), suggesting a possible congenital variation. The possibility of significant contamination of cancer cells in blood samples could be ruled out because, in cases #29 and #31, there were different somatic mutations in *CREBBP* and *TP53* genes, respectively (Fig. 2A), and they were not detected in blood samples from these cases (Table S3). Interestingly, case #34 showed the same frameshift mutation in *NOTCH1* gene as a somatic mutation (Table 2). As for considerably older cattle, the mutation frequency in *TP53* gene was clearly decreased in EBL cases more than 96 months old (Fig. 2A). However, there was no significant difference between ages of EBL cases with *TP53* wildtype gene and those with a mutated gene, because several young EBL cattle were also free of *TP53* mutation (Fig. 2C). However, ages at development of EBL were significantly younger in the cases with high-impact mutations, which exert a large influence on protein translation, such as stop-gained and frameshift mutation, or with biallelic mutations (Fig. 2D,E).

### Analysis of the mutation pattern in SNVs

In all somatic mutations, an advanced analysis focusing on the SNVs was performed. Given the small number of SNVs per EBL case in this study, the 36 EBL cases were grouped by their ages into two categories: an old group over 36 months old (≥ 36-m-old) and a young group less than 36 months old (< 36-m-old). Mutation patterns were then compared among those groups and all cases. The frequency of base substitution from the reference allele to the alternative allele among detected SNVs indicated a large number of mutations involving G-to-A and its complementary sequence C-to-T in EBL cattle (Fig. 3A). That tendency was particularly more pronounced in the ≥ 36-m-old group (G-to-A, 0.38; C-to-T, 0.44) than in the < 36-m-old group (G-to-A, 0.33; C-to-T, 0.25).

**Figure 3.**
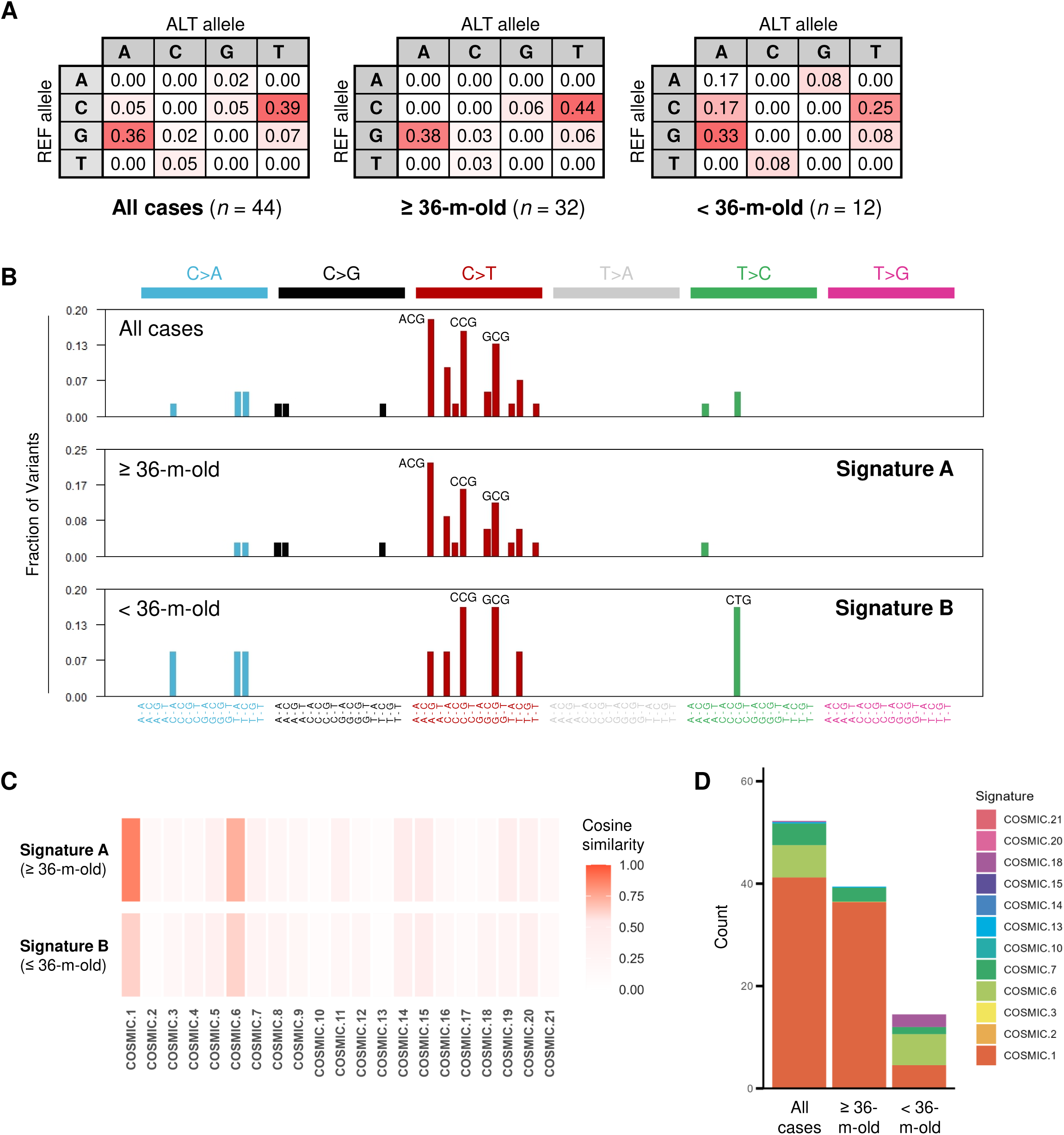
Analysis of mutation patterns in identified single nucleotide variants (SNVs). (A) Rates of nucleotide substitutions in SNVs from 36 EBL cases are shown with a relative color scale. Identified SNVs are put together in all cases or in two groups divided on the basis of cattle age in months. The identical somatic mutation observed in different cases were counted independently. ≥ 36-m-old, group of EBL cases over 36 months old; < 36-m-old, group of EBL cases less than 36 months old. (B) Mutation spectra based on single base substitution (SBS) signatures are drawn using the MutSignatures R package. Identified SNVs are divided into 96 different contexts and their mutation frequencies calculated. The top 3 tri-nucleotide mutations in each group are marked in the spectra. Mutation spectra in the ≥ 36-m-old group and the < 36-m-old group are defined as Signatures A and B, respectively. (C) Heatmap shows similarity between Signature A/B and known COSMIC signature 1–21. Cosine distances between each pair of signatures were calculated and are shown by color intensity. (D) Evaluation of the mutation process using known COSMIC signatures was performed using MutSignatures. The vertical axis indicates the number of mutations imputed for each signature. COSMIC signatures used for estimation are sorted according to validated evidence and reasonable involvement in tumor development.

### Evaluation of single base substitution (SBS) signatures

In the SBS signature, SNVs can be classified into 96 different contexts by considering 6 types of base substitutions (C>A, C>G, C>T, T>A, T>C and T>G, including complementary strands), as well as 4 types of base immediately 5’ and 4 types of base immediately 3’. Mutation spectra were compared based on SBS signatures among all cases, the ≥ 36-m-old group, and the < 36-m-old group (Fig. 3B). As a result, 3 mutation contexts, A[C>T]G, C[C>T]G and G[C>T]G, were found to predominate in all cases and the ≥ 36-m-old group, which indicated that cytosine with guanine on the 3’ side, i.e., cytosine at CpG sites, tended to be converted to thymine. Interestingly, hotspot mutations in *TP53* gene, observed across multiple EBL cases and/or reported in previous studies, corresponded to CpG sites with only one exception (p.Phe229Leu, T[C>A]A) (Table 2). In contrast, of 12 SNVs in *TP53* gene first reported in this study, eleven were inconsistent with CpG sites. The mutation spectrum in the < 36-m-old group showed that the context C[T>C]G, in addition to C[C>T]G and G[C>T]G, was slightly predominant (Fig. 3B), although the small number of SNVs in the young group should be considered.

At this point, the mutation spectrum obtained from SNVs in the ≥ 36-m-old group was defined as Signature A, and that obtained from SNVs in the < 36-m-old group was defined as Signature B. The similarities of mutation context patterns were compared between Signature A or B and the reference signatures, classically known COSMIC signatures 1–21 (Fig. S2A) [15, 16]. Both Signature A and B were similar to COSMIC signatures 1 and 6, and Signature A in particular showed the highest cosine similarity with COSMIC signature 1 (Fig. 3C). We then attempted to evaluate the mutation process of oncogenesis by mutation spectra for all cases, the ≥36-m-old group, and the <36-m-old group using MutSignatures R package [19]. The COSMIC signatures used for the evaluation were selected based on validated evidence and their likelihood of involvement in EBL development (Fig. S2A); for example, COSMIC signature 4 (tobacco smoking) and signature 11 (temozolomide chemotherapy) were excluded because they were less likely to be associated with cattle diseases. As a result, the main components of the mutation process in EBL cases were COSMIC signatures 1, 6, 7, and 18 (Fig. S2B), and above all, COSMIC signature 1 accounted for the majority in all cases and the ≥ 36-m-old group (Fig. 3D). However, not only signature 1, but also COSMIC signature 6 seemed to be largely involved in EBL development in the <36-m-old group. According to the mutation profiles in the COSMIC database, the mutational processes of COSMIC signatures 1 and 6 are “spontaneous deamination of 5-methylcytosine” and “mismatch repair (MMR) deficiency”, respectively (Fig. S2A).

## Discussion

In this study, somatic mutations of 21 cancer-related genes were analyzed in cancer cells from EBL cattle, clearly distinguishing between somatic and germline mutations. The results uncovered several key findings: 1) somatic mutation based on the “two-hit” theory, such as multiple mutations on the same genes or LOH-associated mutations, were frequently observed at EBL onset (Fig. 2A); 2) many EBL cases relied on somatic mutations in *TP53* gene for oncogenesis (Fig. 2B); and 3) CpG sites were the main targets of these somatic mutations (Fig. 3B and Table 2). It is well-known that CpG sites are major targets for methylation induced by cell division, and that 5-methylcytosine is prone to deamination to thymine, leading to a G·T mismatch in double-stranded DNA [20–23]. Thus, the C-to-T mutations at CpG sites are related to aging of the animal as a mutation process, which is supported by the observation that the number of mutations correlates with patients’ ages in COSMIC signature 1 [16]. COSMIC signature 1 is hence considered to be a cell division clock. Besides, an increase in the rate of LOH can result from mitotic recombination [24]. Consequently, the present study suggests that physiological and accidental mutations or chromosomal abnormalities caused by repeated cell division in infected cells for a long period are the factors contributing to EBL onset, particularly in cattle over 36 months old. However, it is important to note that the ages of cattle when EBL developed may not correctly reflect the period of disease progression from viral infection to lymphoma onset, since the time of BLV infection varied among EBL cases.

Regarding the associations between EBL onset in young cattle and somatic mutation patterns, the presence of congenital variations, high-impact mutations significantly affecting protein translation, and biallelic mutations primarily in *TP53*, *NOTCH1*, and *CREBBP* genes may elevate the probability of EBL development in a short period (Fig. 2A,D,E). *TP53* gene is a representative tumor suppressor gene coding p53, which acts as a transcription factor upregulated in response to DNA damage, activating downstream genes involved in cell-cycle control, apoptosis, and DNA repair [25]. NOTCH1 is a receptor protein expressed on the cell membrane that regulates cell differentiation and self-renewal by interaction with ligands on adjacent cells [26]. CREBBP is a lysine acetyltransferase modulating chromatin accessibility through histone acetylation [27]. Although *NOTCH1* and *CREBBP* genes can act as both oncogenes and tumor suppressor genes, they probably functioned as tumor suppressor genes at EBL onset, because most of their mutations were inactivating forms like stop-gained and frameshift mutations (Fig. 2A). For these reasons, somatic mutations in *TP53*, *NOTCH1*, and *CREBBP* genes would result in accelerating the accumulation of secondary mutations due to deficiencies in DNA damage repair and cell division regulation.

Further, the evaluation of the mutation process based on the known signatures showed that COSMIC signature 6, not only signature 1, was mainly involved in oncogenesis in the < 36-m-old group (Fig. 3D). It should be considered that the number of SNVs from the < 36-m-old group was quite small, and hence the accuracy of the estimation by mutation signature analysis might be insufficient; therefore, further analyses are needed to confirm if COSMIC signature 6 is involved in young onset of EBL. Nevertheless, the involvement of COSMIC signature 6, representing MMR deficiency, is worth considering as a possible factor causing young EBL onset, because the evaluation results for SNVs from all cases (*n* = 44) included COSMIC signature 6, whereas that from the ≥ 36-m-old group (*n* = 32) did not (Fig. 3D).

Although the mechanism of EBL development had not been fully elucidated, Rosewick et al. reported that cancer cells from animals developing EBL had proviruses integrated near cancer driver genes, and that these proviruses could increase survival and proliferation in infected cells by disturbing host gene expression [8]. They concluded in their report that it will result in extending the half-life of infected cells and promote the accumulation of further secondary mutations in the genome of oncogenic candidate clones. The present results strongly support this model for EBL development by clarifying the detail of how secondary somatic mutations accumulated in cancer cells. Now, we can propose a renewed hypothesis to explain the general mechanisms of EBL onset; three events need to be completed for tumor development in BLV-infected cattle (Fig. 4). The first event is provirus integration near cancer driver genes, as shown in the previous study [8], which is randomly decided during the infectious cycle in the early phase of BLV infection. The second event is an accidental single mutation (the first “hit”) at CpG sites in cancer-related genes, particularly *TP53*, *NOTCH1*, and *CREBBP* genes, caused by cell division repeated over long periods. The third event is an additional mutation on another allele or chromosomal aberration like LOH as the second “hit”. The oncogenic transformation of BLV-infected cells requires the occurrence of all three random events in the same cell clone, and thus the rate of EBL onset in infected cattle becomes very small.

**Figure 4.**
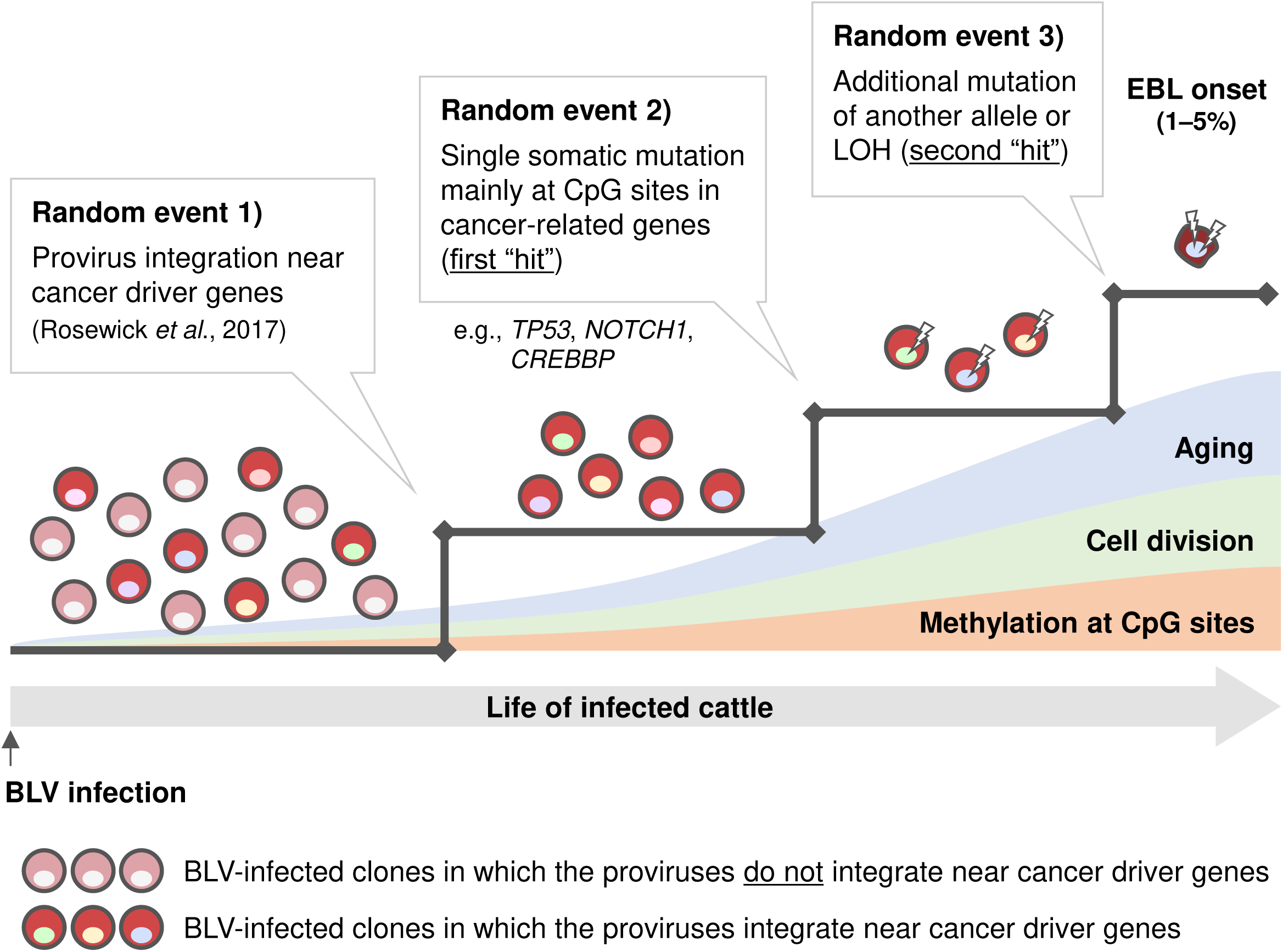
Hypothesis of the general mechanisms of EBL onset. There are 3 events for BLV-infected cells to be transformed into neoplasms. First, infected cell clones whose proviruses are integrated near cancer driver genes acquire long life and reproductive activity [8]. Other clones in which the proviruses do not integrate near cancer driver genes cannot overcome this first obstacle, and thus they are not able to be candidate future cancer cells. Second, single somatic mutation occurs accidentally mainly at CpG sites in cancer-related genes, e.g., *TP53*, *NOTCH1,* and *CREBBP*, as a first “hit”. Finally, an additional mutation or loss of heterozygosity (LOH) occurs on the left normal allele as a second “hit”. These 3 random events need to occur in the same infected clone and be completed during the lives of infected animals, which is considered to be the reason why only a small fraction of BLV-infected cattle develops EBL after a long latent period.

Based on our hypothesis, we can discuss host or viral factors previously reported to be involved in EBL development. The associations between EBL onset and types of cattle major histocompatibility complex BoLA-DRB3 allele [28], viral genotypes [29], or provirus loads during mitotic cycle [30] have been reported in previous studies. At this point, we consider that these factors are related to expanding the diversity of provirus integration sites. The diversity of provirus integration sites, namely clonal variation of infected cells, is increased during the infectious cycle when BLV infects new target cells, followed by a large decrease in clone numbers due to negative selection by the host immune response [7]. In addition, the provirus loads in the chronic phase of BLV infection tend to reflect the number of infected cell clones in the blood [7]. Therefore, these host or viral factors may contribute to a weak immune response against BLV or efficient viral replication in an early phase, resulting in an increase in initial candidate clones with the potential for transformation.

In the research field of ATL caused by HTLV-1 infection, somatic mutation analysis by whole exome sequencing (WES) was conducted using clinical samples from ATL patients. Similar to the present results, age-related C-to-T transitions at CpG sites were predominantly observed on mutation signature analysis [14]. In contrast, the mutation frequency in *TP53* genes of ATL samples was around 20% on WES analysis, suggesting that the oncogenic process in ATL development was less dependent on *TP53* mutations than in EBL cases. In addition, the rate of detecting somatic mutations was reported as 2.3 mutation/Mbp/samples according to the WES in ATL patients. Since the total length of the CDS region in our target resequencing is approximately 73 kbp (Table 1), focusing on selected cancer-related genes worked well for efficiently detecting somatic mutations in the present study. As a limitation of the present study, which is also relevant to WES analysis, a portion of EBL cases (6/36) did not show any somatic mutation in the targeted genes (Fig. 2A), probably because the targeted-gene panel was selected based on information from various human B-cell lymphomas, rather than actual data from WES analysis in cattle. Moreover, the experimental approach cannot identify somatic mutations in promoter regions or epigenetic inactivation. For these reasons, further efforts will be required to improve the methods or study approaches for mutation analysis in bovine B cells.

In this study, several characteristic EBL cases were identified. As for cases #29 and #31, which showed a possible congenital variation in *NOTCH1* genes, they both harbored biallelic and high-impact somatic mutations on *CREBBP* or *TP53* genes, making it unclear whether that congenital variation directly caused the young onset of EBL (Fig. 2A). Moreover, these cases were inconsistent with the characteristics of hereditary cancer, such as Li-Fraumeni syndrome, because there was no second “hit” on *NOTCH1* genes in their genome. However, it is known that *NOTCH1*, as well as *CREBBP*, can be affected by a single mutation due to haploinsufficiency [26, 27, 31]. Based on the observation that case #34 also exhibited the same frameshift mutation as a somatic mutation, the identified congenital variation is likely a functional driver mutation that plays a role in oncogenesis. Unfortunately, there is no additional information on cases #29 and #31 regarding their history, and thus, we cannot discuss their blood relationship. Let us consider two possibilities for the congenital variant; it may be a variant inherited from the parents, or it may have incidentally occurred during the period from fertilized egg to early embryonic development.

For the three cases that did not exhibit biallelic mutations in *TP53* gene, only a single *TP53* mutation was observed in each case (#7, #18, and #28) (Fig. 2A). The somatic mutations from cases #7 and #18, p.Arg167His (R167H) and p.Arg241Trp (R241W), respectively, were mutational hotspots in *TP53* gene, which were also detected in other EBL cases reported in previous studies (Table 2) [9–12]. According to the previous study examining the functions of mutated *TP53*, R241W was not only a loss-of-function mutation impairing the transcriptional activity of p53 protein, but it also acted as a dominant negative variant inhibiting wildtype p53 in the presence of both variants [12]. Conversely, R167H is known as an orthologous mutation of R175H in human *TP53* gene, which is one of the major hotspot mutations observed in various human cancers, showing a dominant negative effect [9, 32–34]. Altogether, it is suggested that the introduction of mutations with dominant negative effects as the first “hit” resulted in oncogenic transformation by inhibiting the normal allele in cases #7 and #18. However, all other cases with R167H or R241W mutations (cases #2, #12, #13, #16, and #19) showed the biallelic *TP53* mutations (Table S3). This suggests that the dominant negative effects of these mutations may not completely suppress the function of the normal allele, and that biallelic inactivity is more advantageous for oncogenesis.

Lastly, regarding the association of age with EBL development, it is interesting that EBL in cattle more than 96 months old showed less dependence on *TP53* mutations (Fig. 2A). This finding may be linked to immunosenescence [35]; the decline in the functions of the immune response due to aging causes a weakened elimination of precancerous abnormal cells, creating a tumor-prone condition. Considering that mutation frequencies of *TP53* gene in diffuse large B-cell lymphoma, one of the nonviral and sporadic human cancers, range from 17.6% to 23.2% [36], the development of EBL in very old cattle may resemble that of spontaneous tumors rather than virus-associated tumors.

In conclusion, this study provides novel insights into the pathogenic mechanism of somatic mutations in cancer cells for EBL development. Furthermore, the present results suggest the possibility that the mutation patterns in young cattle developing EBL are different from those in EBL cattle at common ages. To strengthen the hypothesis inferred from our results, it is necessary to accumulate data using a larger number of clinical samples in the future.

## Materials and methods

### Study design

Fig. 1 summarizes the study strategy. Two samples each were collected from 36 cattle with EBL; one was a blood sample mainly containing normal cells, and the other was a tumor sample mainly containing cancer cells. Because it is known that several EBL cases do not show visible lymphocytosis when a tumor develops and thus, cancer cells are hardly detected in the peripheral blood [17, 37], such EBL cases not showing leukemic signs were selectively used. DNA was extracted from blood and tumor samples and used for PCR amplification of 21 targeted cancer-related genes, followed by amplicon sequencing and variant calling using next-generation sequencing technology. Two variant lists of blood and tumor samples from the same EBL case were compared, and the variants that existed in only tumor samples were identified as somatic mutations. All identified somatic mutations were used for mutation pattern analysis.

### Clinical samples from EBL cattle

The information on EBL cases used in this study are listed in Table S1. All cases were diagnosed as EBL in a domestic livestock hygiene service center and meat inspection center in Japan, and they were kindly provided for research use in the previous studies [37, 38]. To select EBL cases not showing leukemic signs, it was confirmed that lymphocyte counts or clonality values measured by the RAISING-CLOVA method [38] were low in the blood (Table S1). In cases #1–32, peripheral blood mononuclear cells (PBMCs) were isolated from blood samples using Percoll (GE Healthcare Life Sciences, Little Chalfont, UK) density gradient centrifugation for DNA extraction. In cases #33–36, DNA was extracted directly from whole blood samples. Cancer cells were isolated from tumor samples, as described previously [37]; briefly, tumor samples were cut into small pieces, suspended in phosphate-buffered saline, and passed through a 40-μm cell strainer (Corning Inc., Corning, NY, USA) for removal of cell debris. For DNA extraction, the Dneasy Blood and Tissue kit (QIAGEN, Hilden, Germany), Wizard Genomic DNA purification kits (Promega, Madison, WI, USA), or Quick-DNA Miniprep Kits (Zymo Research, Irvine, CA, USA) were used.

### Amplicon sequencing of cancer-related genes

To determine the targeted cancer-related genes, the COSMIC (Catalogue Of Somatic Mutations In Cancer) [39] Cancer Browser database (https://cancer.sanger.ac.uk/cosmic/browse/tissue) was used, and 21 genes that showed high mutation frequencies in various human B-cell lymphomas were selected (Table 1). Primer sets that included all CDS of the 21 targeted genes were designed for long PCR, and 3–4 sets each were mixed for amplification as multiplex PCR (multiplex set A–U, Table S2). The targeted genes in the DNA sample from EBL cattle were amplified using the multiplex primers in a reaction mixture containing 12.5 µl of KOD One PCR Master Mix (TOYOBO, Osaka, Japan), 3.0 µl of primer mix (each 2.5 µM), 0.5 µl of dimethyl sulfoxide (FUJIFILM Wako, Osaka, Japan), 0.2–1.0 µl of DNA template, and extra double-distilled water up to 25 µl. The PCR conditions were as follows: a two-step procedure consisting of 10 sec at 98 °C and 2 min at 68 °C for 25–35 cycles decided depending on the concentration of the template DNA. Following DNA purification using the QIAquick PCR Purification Kit (QIAGEN), the amplicons were pooled and submitted to Macrogen Japan Co., Ltd. (https://www.macrogen-japan.co.jp/) for the DNA-seq analysis. Briefly, a sequence library was prepared using the TruSeq Nano DNA Kit (Illumina, San Diego, CA, USA), and next-generation sequencing analysis was conducted using a NovaSeq 6000 (150-bp pair-end reads, approximately 5.7-Gbp total read bases per sample).

### Data analysis

The identification of somatic mutations was performed using “Variant Calling Pipeline using GATK4” in accordance with online instructions (https://github.com/gencorefacility/variant-calling-pipeline-gatk4) with some modifications. The FASTQ data obtained in DNA-seq of blood and tumor samples from 36 EBL cattle are deposited in the DDBJ database as submission no. DRR531829– DRR531900. The reads were trimmed using Trimmomatic version 0.39 and mapped against the reference genome (*Bos taurus*: ARS-UCD1.2 also known as GCF_002263795.1) using the MEM algorithm in BWA version 0.7.17. The output BAM files were indexed using samtools version 1.6. The variant calling was then conducted for each BAM file using HaplotypeCaller in GATK version 4.3.0.0, and base quality scores were adjusted by GATK’s BQSR algorithm. The variant filtering was performed using GATK’s VariantFiltration with the following criteria: QD < 2, FS > 60, MQ < 40, QUAL < 30, SOR > 4, MQRankSum < −12.5, ReadPosRankSum < −8 for SNVs, and QD < 2, FS > 200, QUAL < 30, SOR > 10, ReadPosRankSum < −20 for insertions or deletions (INDELs). Gene annotations of called variants were added in VCF files using SnpEff version 5.1 with the reference genome, ARS-UCD1.2.99. The variants called only in tumor samples were sorted using bcftools version 1.9.

Of the somatic mutations called by the above procedure, the variants whose read depth was less than 20 or which agreed with known Variant ID in the Ensembl database were excluded from the following analysis. When somatic mutations were also called in blood samples, both VAFs of blood and tumor samples were compared statistically, and it was found that VAF was significantly higher in tumor samples than in blood samples. The definition of chromosome zygosity at the detected somatic mutations was as follows: VAF > 0.7 was considered homozygosity, VAF 0.3–0.7 (SNVs) or VAF 0.2–0.7 (INDELs) was considered heterozygosity, and VAF < 0.3 (SNVs) or VAF < 0.2 (INDELs) was considered a minor mutation. Fig. S1 indicates the interpretation and classification of the patterns of somatic mutations. In comparison with a single heterozygous mutation, multiple heterozygous mutations on a certain target gene were interpreted as meaning that one mutation existed on one allele, and a second mutation existed on a different allele of a homologous chromosome. In addition, a single homozygous mutation was interpreted as a mutation with LOH that resulted in loss of one copy of the target gene by chromosomal abnormalities such as recombination and deficiency. All somatic mutations identified in this study were visually checked in the BAM files using IGV software [40].

### Mutation signature analysis

R package MutSignatures version 2.1.5 [19] with R version 4.3.0 was used for mutation signature analysis of EBL cattle using VCF files generated above in accordance with online instructions (https://cran.r-project.org/web/packages/mutSignatures/vignettes/get_sarted_with_mutS ignatures.html). Bovine whole genome sequences, BSgenome.Btaurus.UCSC.bosTau9, were used as a reference genome. The reference signature datasets listed on the COSMIC website [39] were obtained by the getCosmicSignatures command.

### Statistical analysis

Fisher’s exact test was used to evaluate differences between VAFs of blood and tumor samples when somatic mutations were called in both samples. Associations between cattle ages when EBL developed and patterns of somatic mutations in cancer cells were explored using Mann-Whitney’s U test. A p-value less than 0.05 was defined as significant.

## Appendix

The FASTQ data in this study were deposited in the DDBJ database as submission no. DRR531829–DRR531900. All supplemental information is shown in the supplemental material.

## Acknowledgements

The authors are grateful to the staff at the livestock hygiene service centers and meat hygiene inspection centers in Hokkaido, Iwate, Ibaraki, and Hyogo prefectures in Japan for kindly providing clinical EBL samples. Xiao Bo provided excellent technical support.

This study was supported by JSPS KAKENHI Grant-in-Aid for Early-Career Scientists (19K16008 and 23K14101) (A.N.). The funder had no role in study design, data collection and interpretation, or the decision to submit the work for publication.

A.N. designed the research. A.N., K.A., and T.O. performed data collection and analysis. K.A., Y.M., T.O., and S.K. provided intellectual input, laboratory materials, and/or analytic tools. A.N. drafted the manuscript. All authors read and approved the manuscript and contributed to manuscript revisions.

---

**Supplemental figure 1** Diagram of data interpretation of somatic mutation analysis in this study. Images of the cording sequence (CDS) of targeted genes, read coverage in the IGV view, and homologous chromosomes are shown. (A) In normal cells, there are only germline mutations indicated in black on the targeted gene. (B) Single heterozygous somatic mutation indicated in red is observed in the CDS region of cancer cells. It is interpreted to be located on 1 allele of the homologous chromosome (pink). (C) Double somatic mutations are heterozygously observed in the same targeted gene. This result is interpreted to mean that the first mutation is located on one allele (pink), with the second mutation on another allele (sky blue). (D) Single homozygous somatic mutation is observed, which is interpreted as meaning that, following single heterozygous mutation on one allele, loss of heterozygosity (LOH) occurs by chromosomal recombination or deficiency.

**Supplemental figure 2** Information on known COSMIC signatures obtained from the database. (A) In COSMIC single base substitution (SBS) signatures 1–21, mutational process, supported evidence, and presence of validation for real signatures are listed based on the COSMIC website. (B) Mutation spectra of COSMIC SBS signatures 1, 6, 7 and 18 are drawn using MutSignatures R package and reference signatures datasets. Tri-nucleotide mutations at notable peaks in each signature are marked.

**Supplemental Table 1. Basic information on clinical EBL cases evaluated in this study**

**Supplemental Table 2. Primers used for amplicon sequencing**

**Supplemental Table 3. Details of somatic mutations identified in each EBL case**

**Supplemental Table 4. Loss of heterozygosity of germline mutations in chromosome 19**

