## Supplemental figures for "Effect of C-to-T transition at CpG sites on tumor suppressor genes in tumor development in cattle evaluated by somatic mutation analysis in enzootic bovine leukosis"

**Figure S1**

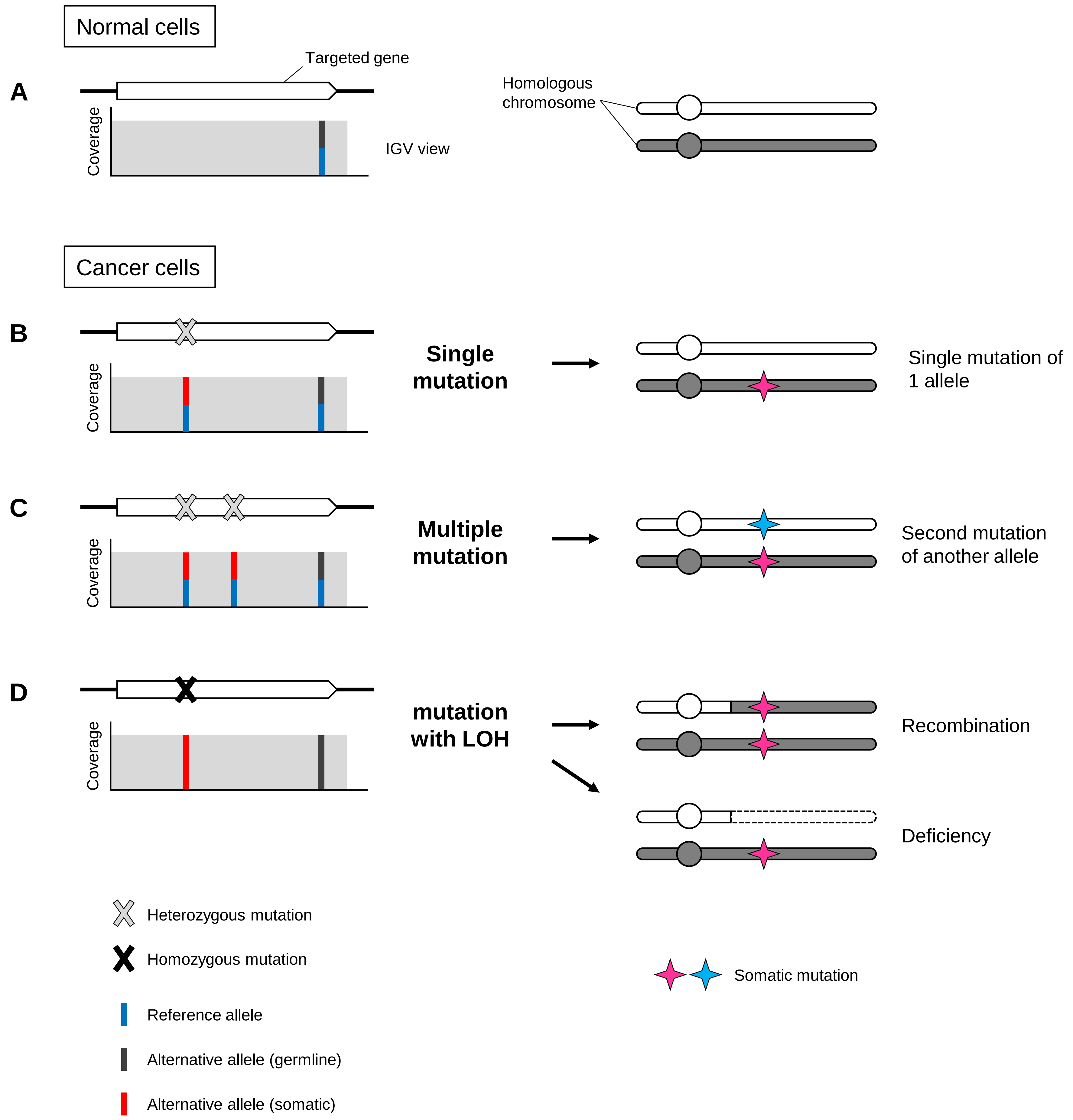
Normal cells
A
Targeted gene
Coverage
IGV view
Homologous
chromosome
Cancer cells
B
Coverage
Single
mutation
Single mutation of
1 allele
C
Coverage
Multiple
mutation
Second mutation
of another allele
D
Coverage
mutation
with LOH
Recombination
Deficiency
Heterozygous mutation
Homozygous mutation
Reference allele
Alternative allele (germline)
Alternative allele (somatic)
Somatic mutation

### Figure S2

A

| COSMIC signature | Proposed etiology | | Validated evidence for real signature? |
|---|---|---|---|
| | Mutational process | Supported by | |
| 1 | Spontaneous deamination of 5-methylcytosine (Aging) | mutational pattern | yes |
| 2 | AID/APOBEC activity | experimental confirmation | yes |
| 3 | HR deficiency | experimental confirmation | yes |
| 4 | Tobacco smoking | experimental confirmation | yes |
| 5 | Unknown (Aging / Tobacco smoking / NER deficiency) | age correlation / statistical association | no |
| 6 | MMR deficiency | statistical association; experimental studies | yes |
| 7 | UV light exposure | experimental confirmation | yes |
| 8 | HR deficiency / NER deficiency | statistical association | no |
| 9 | Polymerase eta somatic hypermutation | statistical association | no |
| 10 | POLE exonuclease domain mutation | experimental confirmation | yes |
| 11 | Temozolomide chemotherapy / MMR deficiency + temozolomide | experimental studies | no |
| 12 | Unknown | unknown | no |
| 13 | AID/APOBEC activity | experimental confirmation | yes |
| 14 | MMR deficiency + POLE mutation | experimental confirmation | yes |
| 15 | MMR deficiency | statistical association; experimental studies | yes |
| 16 | Unknown | unknown | no |
| 17 | Damage by ROS | statistical association | no |
| 18 | Damage by ROS | experimental confirmation | yes |
| 19 | Unknown | unknown | no |
| 20 | MMR deficiency + POLD1 mutation | statistical association; experimental studies | yes |
| 21 | MMR deficiency | statistical association; experimental studies | yes |

HR, homologous recombination; NER, nucleotide excision repair; MMR, mismatch repair; POLE, polymerase epsilon

B

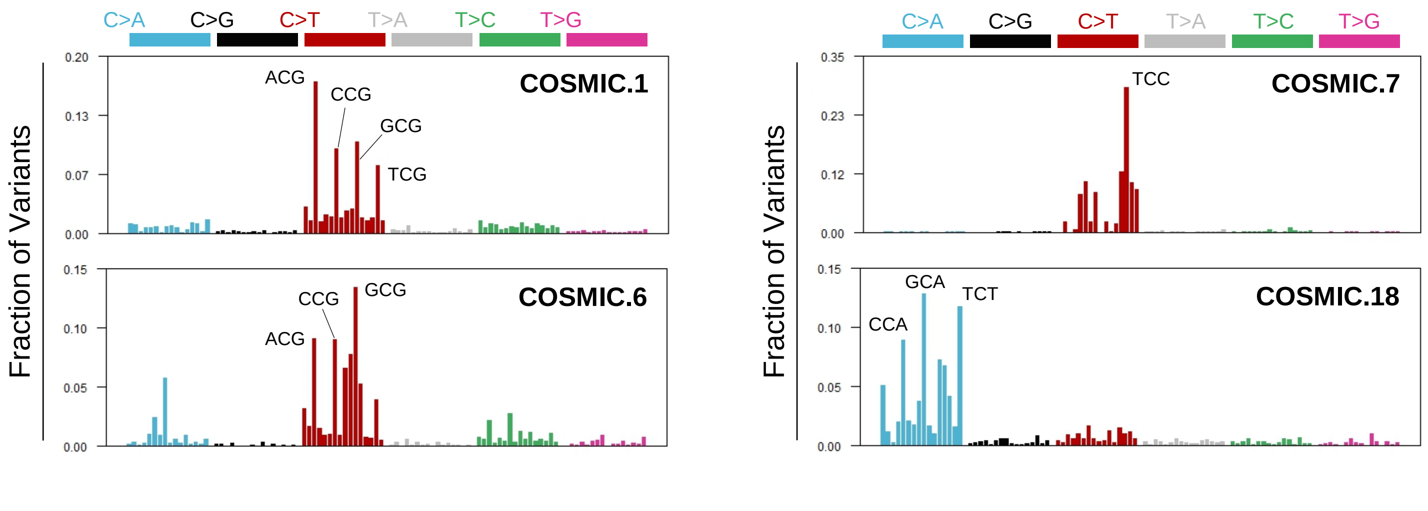
