## Supplemental table 1 for "Effect of C-to-T transition at CpG sites on tumor suppressor genes in tumor development in cattle evaluated by somatic mutation analysis in enzootic bovine leukosis"

### Supplemental Table 1. Basic information on clinical EBL cases evaluated in this study

| Case No. | ID | Age (months) | Sex[1)] | Breed[2)] | WBC (cells/µl) | Lym (cells/µl) | Cv blood | Cv tumor |
|---|---|---|---|---|---|---|---|---|
| 1 | 28-18 | 31 | F | JB | 13,900 | 6,000 | - | - |
| 2 | 24-14 | 43 | F | HO | 6,000 | 1,500 | - | - |
| 3 | 10-22 | 74 | F | HO | 13,100 | 7,900 | - | - |
| 4 | 45-7 | 67 | F | HO | 9,600 | 3,300 | - | - |
| 5 | 26-23 | 50 | F | HO | 10,200 | 4,700 | - | - |
| 6 | 29-36 | 67 | F | HO | 13,400 | 4,800 | - | - |
| 7 | 33-4 | 88 | F | HO | 6,700 | 2,200 | - | - |
| 8 | 53-10 | 181 | F | JB | 8,100 | 4,700 | - | - |
| 9 | 21-35 | 91 | F | HO | 13,100 | 3,400 | - | - |
| 10 | 27-24 | 25 | M | JB | 5,100 | 2,600 | - | - |
| 11 | 38-44 | 28 | F | JB | 4,600 | 1,800 | - | - |
| 12 | 40-6 | 81 | F | HO | 4,900 | 1,700 | - | - |
| 13 | 41-42 | 82 | F | HO | 7,200 | 2,700 | - | - |
| 14 | 44-39 | 82 | F | HO | 9,690 | 5,460 | - | - |
| 15 | 46-20 | 48 | F | HO | 25,800 | 2,000 | - | - |
| 16 | 50-9 | 83 | F | HO | 7,600 | 3,800 | - | - |
| 17 | 52-40 | 75 | F | HO | 8,300 | 1,770 | - | - |
| 18 | 34-38 | 84 | F | HO | 15,400 | 8,400 | - | - |
| 19 | 18-11 | 133 | F | JB | 30,700 | 9,800 | - | - |
| 20 | 20-17 | 127 | F | JB | 1,600 | 1,300 | - | - |
| 21 | 22-3 | 54 | F | HO | 5,000 | 2,100 | - | - |
| 22 | 30-19 | 78 | F | HO | 5,200 | 3,700 | - | - |
| 23 | 43-15 | 111 | F | HO | 41,500 | 12,100 | - | - |
| 24 | 32-37 | 118 | F | HO | 9,400 | 6,700 | - | - |
| 25 | 36-5 | 30 | F | JB | 6,700 | 2,400 | - | - |
| 26 | 39-41 | 136 | F | JB | 8,700 | 3,000 | - | - |
| 27 | 49-43 | 21 | F | JB | 5,130 | 1,670 | - | - |
| 28 | EBL002 | 34 | F | HO | - | - | 0.09 | 1.00 |
| 29 | EBL024 | 17 | F | JB | - | - | 0.30 | 0.82 |
| 30 | EBL065 | 27 | F | F1 | - | - | 0.11 | 0.52 |
| 31 | EBL070 | 29 | M | JB | - | - | 0.39 | 1.00 |
| 32 | EBL098 | 22 | F | HO | - | - | 0.10 | 0.79 |
| 33 | EBL221 | 28 | M | JB | - | - | 0.55 | 0.78 |
| 34 | EBL188 | 26 | M | JB | - | - | 0.31 | 0.95 |
| 35 | EBL119 | 133 | F | HO | - | - | 0.08 | 0.74 |
| 36 | EBL184 | 93 | F | HO | - | - | 0.11 | 1.00 |

1) M, male; F, female

2) HO, Holstein Friesian (dairy cattle); JB, Japanese Black (beef cattle); F1, mix breed (HO×JB)

WBC, whole blood cells; Lym, lymphocytes; Cv, clonality value measured by BLV RAISING-CLOVA
