## Supplemental table 2 for "Effect of C-to-T transition at CpG sites on tumor suppressor genes in tumor development in cattle evaluated by somatic mutation analysis in enzootic bovine leukosis"

### Supplemental Table 2. Primers used for amplicon sequencing

| Gene | | | Location | Sequences (5' → 3') | Product size | Multiplex set |
|---|---|---|---|---|---|---|
| *TBL1XR1* | 1 | F | 90115086 | ATGCTCAAGGCATTGTTCTTAGCTCTGT | 8241 | J |
| | | R | 90123327 | CCACCAATAAAGAATGGAGCTGTTGCAC | | |
| | 2 | F | 90159639 | TTCAGCTGGAGGGATCTTGAATTGGTTC | 11823 | K |
| | | R | 90171462 | TCTCCCCACTTCAACCAAGACCTAGAAT | | |
| | 3 | F | 90171723 | CAAGGAAAGCTGATGACAGACCAGAGAG | 11914 | L |
| | | R | 90183637 | AACACTGTGCCACTGAGTGTAAAAAGGA | | |
| | 4 | F | 90191873 | AGACCCAGTCTTAAAGAGAGAGGACCAC | 11275 | R |
| | | R | 90203148 | CACCCTGAAAGCCTCTTGGTATCACATT | | |
| | 5 | F | 90201776 | TCCATGCATCTGTGGTTTCAGCAGTATT | 8321 | N |
| | | R | 90210097 | CAGTATCTCTTAAAGCAGCACGTTCGGA | | |
| *NRAS* | 1 | F | 28614522 | TGTTCGACTTTTTATCACGGGAACGGAT | 11053 | G |
| | | R | 28625575 | AATGAGGAAACTACTGGCTAGAGGAGCA | | |
| *EZH2* | 1 | F | 112078080 | GCCACTCAGGGAGAAAAATACCTTCCAA | 8267 | F |
| | | R | 112086347 | AAACAAGCCTCATCAAAACCTCCATCCT | | |
| | 2 | F | 112059203 | GAAATGGAGAACCCCAGAAGAAGTCAGG | 11234 | G |
| | | R | 112070437 | TACCTTCACAGGTGATTGTGATGCTTCC | | |
| | 3 | F | 112049746 | TAACACGTGAAGGGCAATGTCAGGATAC | 9484 | A |
| | | R | 112059230 | CCTGACTTCTTCTGGGGTTCTCCATTTC | | |
| | 4 | F | 112038557 | AAACACCAGACTTAACGCTCAACAGACA | 11216 | I |
| | | R | 112049773 | GTATCCTGACATTGCCCTTCACGTGTTA | | |
| | 5 | F | 112028761 | CTAATTCAGAGTCTGGACGACCAAGCAG | 10346 | K |
| | | R | 112039107 | TCGTTTCATACGCTTTTCTGTAGGCGAT | | |
| | 6 | F | 112023097 | CATCTCCTGATGTGGTCACTCATTCCAC | 5534 | U |
| | | R | 112028631 | GTAACTGAGATGGCACTGTCGTCAGAAA | | |
| *KMT2D* | 1 | F | 30757566 | AGTGCAGAGCATATTACAGTGTTTGGCT | 9893 | L |
| | | R | 30767459 | CTGCAGACATAACACACTCTGCTCCTAC | | |
| | 2 | F | 30761730 | CCTTTCACCTCTCCAAGACTCCATTTCC | 10586 | N |
| | | R | 30772316 | ATAACTGATCCCATTCCCCAGACCAGAT | | |
| | 3 | F | 30771219 | TAAACACCCTTTAAATCCTGCCAGTCCC | 11557 | O |
| | | R | 30782776 | AGGACTCTGCTAATCAGACTTCAGGTGT | | |
| | 4 | F | 30781488 | GGGACAAGAAGGACATCTTCAATGAGCA | 11126 | P |
| | | R | 30792614 | TCTGGAGAACTTCAAGAGACCTCTACCG | | |
| | 5 | F | 30790054 | CCAGAGAGCAAACCTTATGGAGTCTTGG | 10550 | Q |
| | | R | 30800604 | CTGGGTGACCAGCTGCAAATCATTTTAC | | |
| *KRAS* | 1 | F | 84747210 | CCAGGAACCAGCATGTATCTTTGGAAGT | 7901 | F |
| | | R | 84755111 | AGTATGCTGGCTTTAGGAACAGGTGAAC | | |
| | 2 | F | 84771071 | TTGGCAAGTGAAGACTGTGAGGAAAAGT | 11777 | T |
| | | R | 84782848 | CACCTGGGTAAAGAAGTGATGCTGATGT | | |
| | 3 | F | 84782821 | ACATCAGCATCACTTCTTTACCCAGGTG | 11294 | I |
| | | R | 84794115 | TTTAGGCTGTGATGGTGCCTTTAAGGAG | | |
| | 1 | F | 112279875 | CCATCAATTTACCCAAGGCCAAACACAG | 3512 | M |
| | | R | 112283387 | TCGTCTTCATTATCCTGGCACCTCAAAC | | |
| | 2 | F | 112299341 | CAGGAGAGGAGGAAGAGAAAGAGACACA | 8186 | N |
| | | R | 112307527 | TGAATATGGCAGGACCCCAAAACTTTCA | | |
| | 3 | F | 112306698 | TGGAGATGAATGGCAAAACCTGCTTAGT | 11112 | O |
| | | R | 112317810 | TTCATTCCATAAACACGCACTTAGGGCT | | |

| | | | | | | |
|---|---|---|---|---|---|---|
| *EP300* | 4 | F | 112313499 | TAATCTTGGCTTGTGCCCTTTGTCAGAT | 11004 | P |
| | | R | 112324503 | TTGCTTGCCTCATTACCATTCAGTAGCA | | |
| | 5 | F | 112323546 | CTTCACAGGAAGCAAGAATGGAGCCTAA | 9050 | Q |
| | | R | 112332596 | ACAATAAAGGAGGCGATTACGAAACGGT | | |
| | 6 | F | 112330407 | GCTCAGGCTTCTAAGTGGTCTTGGATTT | 9085 | R |
| | | R | 112339492 | CTTTTCATTAGGCAACAGCAGCCTTAGC | | |
| | 7 | F | 112338018 | GCCAGAAATACTGACTTCAGACCAGACC | 9705 | S |
| | | R | 112347723 | GCTGGTTTTGAGGATTCAGTGAGCTAGT | | |
| *TET2* | 1 | F | 20036193 | ACTCTGGAAGTCTGTCATAGCAACTCCT | 4260 | H |
| | | R | 20040453 | CGAGAGTCCGTCATTCGGAGTTTAGTTC | | |
| | 2 | F | 19994792 | ACTGAGACAAAGTAGCCAGAGGTGAGAT | 3552 | M |
| | | R | 19998344 | CTCTTTGATGATTTGCCAGCTTGGTTCC | | |
| | 3 | F | 19931817 | ACCAACACCAGCTCGAGATGTCTTAATG | 10427 | E |
| | | R | 19942244 | CTATTAGGCAGGAATTCGCAGAAGGAGG | | |
| | 4 | F | 19913146 | AAGAAACACCATGCAGGAGTCTGTGAAT | 9347 | F |
| | | R | 19922493 | TAAGACTGTTGTTGTCCAGTGAGGTCAG | | |
| | 5 | F | 19894676 | GTTACTCCATTCCTGACACTGGCTACAC | 11429 | G |
| | | R | 19906105 | AAAGCATTTGCCCCATTTCACCTCATTC | | |
| *TNFAIP3* | 1 | F | 75739861 | GGGGACACCAAGGCAATAAAAGGACTTA | 10824 | O |
| | | R | 75750685 | CTTGAACGGGGATTTCTACCACCATCAA | | |
| | 2 | F | 75748250 | AGCAGAGAGGCTGGTTTATTCTGGAAAC | 9435 | A |
| | | R | 75757685 | TTGGCAAGGTGTTGATTGTTGAAACTGG | | |
| *B2M* | 1 | F | 103094337 | CCACCAGGTAACGTCAGCTCCTTTTTAT | 8755 | B |
| | | R | 103103092 | AGATCACAGCACCACCAAACTTATCTAAC | | |
| | 2 | F | 103102027 | GGTATTTCTTCACAGGCTCTTCTGCCAT | 8937 | C |
| | | R | 103110964 | AGCAGCCACCTAAGATGTTCATTCTCAC | | |
| *NOTCH1* | 1 | F | 103976741 | GAAACAAGATTTAGGGCATCAAGCGTCG | 5409 | H |
| | | R | 103982150 | AGGGGCCTAATTGGTGATTGGAAAAACT | | |
| | 2 | F | 103952668 | TTTAGATAGAGCCAATGCCAGGCACGTA | 8892 | B |
| | | R | 103961560 | GTAGGCAGGTCCCTGAAACAAAAGGTTG | | |
| | 3 | F | 103945505 | AACACTTGTAGGTGTTGGTGAGGTCGAT | 8879 | C |
| | | R | 103954384 | CTTGCAGATCAGGGTCTCAGGACTATGG | | |
| | 4 | F | 103937522 | GGTTTTCTCATCTGTACAGCGGACATGC | 10545 | E |
| | | R | 103948067 | GTCCATTCTCCCTTGGGGATTAGAAGCA | | |
| | 5 | F | 103933639 | AGAACTTACCCCACTCACTATGTCACCA | 9045 | D |
| | | R | 103942684 | CAAGATGGGGAAAGAGACCTGGGATTTC | | |
| *ATM* | 1 | F | 17842528 | CCACTGAAGAAGATTCACCGCTAGTCTG | 10417 | L |
| | | R | 17852945 | ATTTTTCTCCCCCTGCAATACCTCACAG | | |
| | 2 | F | 17851849 | TAGGTACCTGCCTGTATGGTTCTTTGGA | 9895 | D |
| | | R | 17861744 | TTCAGAGAACGTGCCAGATGATGGAATC | | |
| | 3 | F | 17864052 | CACAGTCTTGGGGTTTATGGTGATGAGG | 7241 | N |
| | | R | 17871293 | AAACTACTGAGTGGGAGTTTGTAAGGACA | | |
| | 4 | F | 17877642 | ATGCCCTTCTACTTATCTCTGCTTGACC | 7116 | O |
| | | R | 17884758 | ATGACTTCTCTCCCTTTTCATGCAACCA | | |
| | 5 | F | 17888675 | TTGGCCATCAGGACATGCTTCTGTATTT | 9352 | P |
| | | R | 17898027 | AGTGAGCTATGACTGTGCTAGACCTTGA | | |
| | 6 | F | 17897344 | AACATGGTTAGCTGAAGTGTGATAGCCC | 9826 | Q |
| | | R | 17907170 | AGTGTAAGCTGGAAGATTGGGTGAAAGG | | |
| | 7 | F | 17908087 | TGCCTGTAGCAAGAAGTGGGTAATGAAG | 10181 | R |
| | | R | 17918268 | AGCACCTCACATTTCCCCAGTATTTCAG | | |
| | 8 | F | 17915427 | GTTCCTCAGTGTCCATCCCACAGTAAAG | 7527 | S |

| | | | | | | |
|---|---|---|---|---|---|---|
| *ATM* | 8 | R | 17922954 | TTTATTCCCCAAGAGAAGGAGAGGGTGT | 7527 | S |
| | 9 | F | 17923784 | TGGGACTTTTATATGGCAAGTGGCTTGT | 9517 | T |
| | | R | 17933301 | CTTCCTTTTGTCTGGGATTGTCTCCCTC | | |
| | 10 | F | 17931396 | GATCCACAACCCCTGCAAATTTGGATTC | 9577 | J |
| | | R | 17940973 | ACCTGAATTATTCCTGCCTGACGAGTTG | | |
| | 11 | F | 17939736 | GTCTAGCCCTCACCTTTCATTCAACCTC | 8088 | A |
| | | R | 17947824 | AAGATCTCCAAAATGACTGTGCGTAGGG | | |
| | 12 | F | 17945469 | TAGAGATCCTGCAGGCCCTAAAATCCAT | 8065 | B |
| | | R | 17953534 | CCACTTTCTGGAGCGGAACAGTAATACA | | |
| | 13 | F | 17960654 | CCACTGCTGCATCTAGCACAGTAAAATG | 7533 | C |
| | | R | 17968187 | CAGGTACTGCCCATAATCCCATCATACC | | |
| | 14 | F | 17974174 | AAAGGGAGTTTTGTCTCTTTCCCAGCAA | 7154 | D |
| | | R | 17981328 | TAAATCTTCCTGGCCCCTTGTTCCTCTA | | |
| | 15 | F | 17986175 | AAGAAGGCAGGCAGAGCATAGAACAAAT | 11398 | E |
| | | R | 17997573 | GTCACTCAAGATTCCCATGAGAAGGTCC | | |
| *CD79A* | 1 | F | 51344192 | ACTCGCTCTGTCTCTCCATCCTTATCTC | 5520 | B |
| | | R | 51349712 | ACTGACCTGTGGCTGAAATGAAAGAACA | | |
| *TP53* | 1 | F | 27375293 | GGCCCCAGTCTGGCCATCCTTCTAAT | 13285 | T |
| | | R | 27388578 | CTGACTTTCCCCGCACTCTCCTCTCC | | |
| *CD79B* | 1 | F | 48131663 | TAGGTGGAGGACTTTGAAGGACAAGAGG | 4213 | U |
| | | R | 48135876 | AAGGGTATAAGTTGGTCTGGGGTGGAAG | | |
| *MYD88* | 1 | F | 11608904 | CTCCCAATGTCAGTGTTTTCCCCTAGAC | 5211 | M |
| | | R | 11614115 | CCAGGGATCTGTGATCTAGAATGGGGAT | | |
| *PIM1* | 1 | F | 11065620 | AACATAAAAATCTGCCAGGGGATCTGGG | 7639 | A |
| | | R | 11073259 | GGGTAGACATGAGTAGCGGTATTTGCAG | | |
| *BCL2* | 1 | F | 61581094 | CTGCACAGGGTTGGAATTTATGGTCTCT | 10119 | S |
| | | R | 61591213 | GAGTTAAGGCAAGTTCTGAGAAGACCCC | | |
| | 2 | F | 61392362 | CTGAAAGACTCCACACCCTGATCCAATC | 5422 | C |
| | | R | 61397784 | AACTCTGTGCTCTAATTCCAGGTTTGGG | | |
| *CREBBP* | 1 | F | 3168833 | TCAACCATGCTGGAGAATCACACAATCA | 7065 | D |
| | | R | 3175898 | TAAACACCTCCTGCCTTCTTATGCAGTG | | |
| | 2 | F | 3145134 | GAAGCAGTTACCTGTAAGCAGAAGAGGG | 7454 | F |
| | | R | 3152588 | ATGGCGTTAAGATAACTGGTGTCAGTGG | | |
| | 3 | F | 3111634 | AGTGTGTGTACTGGGCACAACTTGTTTA | 11791 | E |
| | | R | 3123425 | CTATACCTAGGGGTGACCCTCATCTGTG | | |
| | 4 | F | 3094675 | CCTCATAAGAAGTCAGTTCGGGGGAAAC | 9963 | G |
| | | R | 3104638 | TACTTCTCCACTCAGTACACTCAGCCTC | | |
| | 5 | F | 3087782 | CCAAGGTGAACCGAAAGAGAGAGGAATC | 7400 | H |
| | | R | 3095182 | GACAGACAGACGGACAGATGGGTATTTG | | |
| | 6 | F | 3076360 | AAGATTTCTCAAGCATCCCTGTCACTCG | 11858 | I |
| | | R | 3088218 | TGTTCCTTTGGATTGTGTTGTGCTTTGG | | |
| | 7 | F | 3064626 | GAACACTCTGACACTGACCTCCAAAACA | 8061 | J |
| | | R | 3072687 | TCATTCGGCTTGTTTACTTCTGCGGTAT | | |
| | 8 | F | 3052806 | ACCTTTTGTTACCCAGATATTGCAGCCA | 11847 | K |
| | | R | 3064653 | TGTTTTGGAGGTCAGTGTCAGAGTGTTC | | |
| *SOCS1* | 1 | F | 9911738 | TTGGAAAACATCACCTCCTCCACTATGC | 3040 | M |
| | | R | 9914778 | TATCCTAATGGGCTTGCCTGAAGAAACG | | |
| | 1 | F | 40415518 | ACCGCAGCATTATGTCTTTCTGCTTAGT | 3963 | U |
| | | R | 40419481 | ATAACAACGGCAAAGGTCGCTGTATGTA | | |
| | 2 | F | 40481333 | ACATCCTGCCTTCTGCACTAATGATGAC | 11727 | P |
| | | R | 40493060 | GGGTTCAGACCTGGAAGACCAGTATTTG | | |

| | | | | | | |
|---|---|---|---|---|---|---|
| *CARD11* | 3 | F | 40492889 | ATGGAAACAGTAAGTGGACCCAGTAGGA | 9829 | Q |
| | | R | 40502718 | GGACAGTCCTGTGGGTTAAAGCAGATAC | | |
| | 4 | F | 40499092 | AGGCTTTCTGAGGACTGTAACTGCTCTA | 9172 | R |
| | | R | 40508264 | GTTACGGATGAGCTGGGTATTCATAGGC | | |
| | 5 | F | 40504522 | ATTCCACAATGGGTATTGAGTGCCTGTT | 11229 | T |
| | | R | 40515751 | ATCGGTGAAGCAGCAGGATTTAAGGAAA | | |
| | 6 | F | 40515724 | TTTCCTTAAATCCTGCTGCTTCACCGAT | 8343 | S |
| | | R | 40524067 | TCCCAGCTCATTAACATACACACAACCC | | |
| *PTEN* | 1 | F | 9462884 | AAGGAAGAGCAGTGCTAATAACCGGAAC | 4532 | H |
| | | R | 9467416 | TTTGCCTTCAAATGAGCACCATACATGC | | |
| | 2 | F | 9492424 | GCTACCCATTTCTCCCTTCTTCAGCTTT | 7004 | J |
| | | R | 9499428 | GAGGCATAAAGGCAGAGGTTTCTGAAGT | | |
| | 3 | F | 9529029 | GTGTTGGCTGTGAGTGTTGTTTAGCTTC | 10501 | I |
| | | R | 9539530 | TAAACATCTCAGGTCCTCTGCTCTGGAA | | |
| | 4 | F | 9550642 | CACGGCTGTGAGAGACAAGAATAGAGTG | 10729 | K |
| | | R | 9561371 | AGGTTTCCTCTGGTCCTGGTATGAAGAA | | |
| | 5 | F | 9555402 | GTCCCTAGCAAACATCTGTCAACTCTCC | 9351 | L |
| | | R | 9564753 | AGCTGGAGATGGTATATGGTCCAGAGTC | | |
