## Supplemental table 3 for "Effect of C-to-T transition at CpG sites on tumor suppressor genes in tumor development in cattle evaluated by somatic mutation analysis in enzootic bovine leukosis"

### Supplemental Table 3. Detail of somatic mutations identified in each EBL case

| Case No. | ID | Chr | Position | REF[1] | ALT[2] | Gene | VAF blood[3] | VAF tumor[3] | Fisher's exact test | Are there Variant IDs at this position? | Remarks |
|---|---|---|---|---|---|---|---|---|---|---|---|
| 2 | 24-14 | 5 | 30793539 | G | A | *KMT2D* | 0.1462 | 0.2945 | $p < 0.0001$ | no | minor mutation |
| | | 19 | 27379331 | C | T | *TP53* | 0.2280 | **0.9878** | $p < 0.0001$ | no | LOH |
| 3 | 10-22 | 19 | 27378376 | C | T | *TP53* | - | **0.9535** | - | no | LOH |
| 4 | 45-7 | 5 | 30792244 | G | T | *KMT2D* | - | 0.5028 | - | rs447261496 (G>C) | - |
| | | 19 | 27380015 | C | T | *TP53* | - | **0.9650** | - | no | LOH |
| 5 | 26-23 | 5 | 84753447 | G | A | *KRAS* | - | 0.3241 | - | no | - |
| | | 25 | 3061108 | G | A | *CREBBP* | - | 0.4867 | - | no | - |
| | | 25 | 3147752 | G | A | *CREBBP* | - | 0.4769 | - | no | - |
| 6 | 29-36 | 19 | 27378819 | C | T | *TP53* | - | **0.9618** | - | no | LOH |
| | | 11 | 103946432 | C | T | *NOTCH1* | - | 0.6622 | - | no | - |
| 7 | 33-4 | 19 | 27379331 | C | T | *TP53* | - | 0.4689 | - | no | - |
| 9 | 21-35 | 19 | 27378769 | AAG | A | *TP53* | - | 0.2428 | - | no | - |
| | | 19 | 27378804 | C | T | *TP53* | - | 0.3233 | - | no | - |
| 10 | 27-24 | 19 | 27378778 | C | A | *TP53* | - | **0.8251** | - | no | LOH |
| 11 | 38-44 | 19 | 27378805 | G | A | *TP53* | - | **0.8661** | - | no | LOH |
| 12 | 40-6 | 19 | 27378808 | G | A | *TP53* | - | **0.8936** | - | no | LOH |
| 13 | 41-42 | 19 | 27379331 | C | T | *TP53* | - | **0.9921** | - | no | LOH |
| 14 | 44-39 | 19 | 27378842 | G | T | *TP53* | - | **0.9713** | - | no | LOH |
| 15 | 46-20 | 19 | 27378804 | C | T | *TP53* | - | **0.9093** | - | no | LOH |
| 16 | 50-9 | 19 | 27379331 | C | T | *TP53* | - | 0.6559 | - | no | LOH* |
| 17 | 52-40 | 19 | 27377694 | G | A | *TP53* | - | 0.4541 | - | no | - |
| | | 19 | 27379338 | C | G | *TP53* | - | 0.4911 | - | no | - |
| 18 | 34-38 | 19 | 27378808 | G | A | *TP53* | - | 0.4743 | - | no | - |
| 19 | 18-11 | 19 | 27379319 | T | C | *TP53* | - | 0.2584 | - | no | minor mutation |
| | | 19 | 27379331 | C | T | *TP53* | - | 0.2223 | - | no | minor mutation |
| 21 | 22-3 | 25 | 3078462 | G | A | *CREBBP* | - | 0.4442 | - | no | - |
| | | 19 | 27378805 | G | A | *TP53* | - | **0.7782** | - | no | LOH |
| | | 26 | 9466850 | G | A | *PTEN* | - | 0.4138 | - | no | - |
| 22 | 30-19 | 5 | 30777496 | G | A | *KMT2D* | - | 0.4460 | - | no | - |
| | | 19 | 27378376 | C | T | *TP53* | - | 0.4553 | - | no | - |
| | | 19 | 27379193 | G | C | *TP53* | - | 0.4060 | - | no | - |
| 23 | 43-15 | 25 | 40519575 | G | A | *CARD11* | - | 0.2381 | - | no | minor mutation |
| 25 | 36-5 | 19 | 27378376 | C | T | *TP53* | - | **0.9996** | - | no | LOH |
| 26 | 39-41 | 5 | 30796297 | C | T | *KMT2D* | - | 0.4813 | - | no | - |
| | | 22 | 11611445 | C | G | *MYD88* | - | 0.5045 | - | no | - |
| 27 | 49-43 | 19 | 27378365 | C | T | *TP53* | - | **0.9765** | - | no | LOH |
| 28 | EBL002 | 19 | 27378780 | A | G | *TP53* | - | 0.4907 | - | rs450926493 (A>C) | - |
| 29 | EBL024 | 25 | 3085919 | T | TG | *CREBBP* | - | **0.8078** | - | no | LOH |
| | | 11 | 103956330 | AC | A | *NOTCH1* | 0.2013 | 0.1327 | $p = 0.0154$ | no | germline mutation |
| 30 | EBL065 | 5 | 84753447 | G | T | *KRAS* | - | 0.3456 | - | no | - |
| | | 19 | 27379449 | G | A | *TP53* | - | **0.7183** | - | no | LOH |
| | | 26 | 9538357 | C | T | *PTEN* | - | 0.1855 | - | no | minor mutation |
| 31 | EBL070 | 19 | 27377679 | G | A | *TP53* | - | **0.7273** | - | no | LOH |
| | | 11 | 103956330 | AC | A | *NOTCH1* | 0.2391 | 0.1644 | $p = 0.0580$ | no | germline mutation |
| 33 | EBL221 | 19 | 27379400 | G | A | *TP53* | 0.2073 | 0.3434 | $p < 0.0001$ | no | - |
| | | 19 | 27380089 | C | A | *TP53* | 0.1486 | 0.2631 | $p < 0.0001$ | rs457482802 (C>T) | minor mutation |
| 34 | EBL188 | 11 | 103956322 | T | C | *NOTCH1* | - | 0.1131 | - | no | minor mutation |
| | | 11 | 103956330 | AC | A | *NOTCH1* | - | 0.1289 | - | no | minor mutation |
| 36 | EBL184 | 19 | 27379974 | C | T | *TP53* | - | **0.8714** | - | no | LOH |

1) REF, reference allele

2) ALT, alternative allele

3) VAF, variant allele frequency; VAF more than 0.70 are shown in bold, and VAF from 0.30 (SNVs) or 0.20 (INDELs) to 0.70 are underlined.

* Although the tumor VAF is less than 0.70, this variant was exceptionally considered to be biallelic because the presence of LOH in a wide range of *TP53* gene was suggested in this case (Table S4).
