## Supplemental table 4 for "Effect of C-to-T transition at CpG sites on tumor suppressor genes in tumor development in cattle evaluated by somatic mutation analysis in enzootic bovine leukosis"

**Supplemental Table 4. Loss of heterozygosity of germline mutations in chromosome 19**

| Case No. | ID | Chr | Position | REF[1)] | ALT[2)] | Gene | VAF blood[3)] | VAF tumor[3)] | Variant ID |
|---|---|---|---|---|---|---|---|---|---|
| 3 | 10-22 | 19 | 48132610 | C | T | *CD79B* | 0.5129 | **0.9710** | rs134114145 |
| | | 19 | 48133157 | G | A | *CD79B* | 0.4837 | - | rs210668899 |
| 6 | 29-36 | 19 | 48132610 | C | T | *CD79B* | 0.5281 | **0.9822** | rs134114145 |
| 11 | 38-44 | 19 | 48132610 | C | T | *CD79B* | 0.4832 | - | rs134114145 |
| | | 19 | 27379196 | C | T | *TP53* | 0.5289 | **0.9257** | rs133909661 |
| 15 | 46-20 | 19 | 48132610 | C | T | *CD79B* | 0.5064 | **0.9614** | rs134114145 |
| | | 19 | 27377007 | A | G | *TP53* | 0.4822 | **0.9503** | rs209064154 |
| | | 19 | 27379184 | C | T | *TP53* | 0.5072 | **0.9576** | rs456002482 |
| | | 19 | 27380071 | C | G | *TP53* | 0.5010 | **0.9621** | rs478425409 |
| 16 | 50-9 | 19 | 27377007 | A | G | *TP53* | 0.5046 | **0.8147** | rs209064154 |
| | | 19 | 27379184 | C | T | *TP53* | 0.5075 | **0.8353** | rs456002482 |
| | | 19 | 27380071 | C | G | *TP53* | 0.5012 | **0.8217** | rs478425409 |
| 22 | 30-19 | 19 | 48132610 | C | T | *CD79B* | 0.4993 | - | rs134114145 |
| 24 | 32-37 | 19 | 48132610 | C | T | *CD79B* | 0.5114 | **0.9833** | rs134114145 |
| 27 | 49-43 | 19 | 27379196 | C | T | *TP53* | 0.4761 | **0.9873** | rs133909661 |
| 31 | EBL070 | 19 | 48133157 | G | A | *CD79B* | 0.4223 | 0.1936 | rs210668899 |
| 36 | EBL184 | 19 | 27377007 | A | G | *TP53* | 0.5937 | **1.0000** | rs209064154 |
| | | 19 | 27379184 | C | T | *TP53* | 0.5352 | **1.0000** | rs456002482 |
| | | 19 | 27380071 | C | G | *TP53* | 0.5518 | **1.0000** | rs478425409 |

1) REF, reference allele

2) ALT, alternative allele

3) VAF, variant allele frequency; VAF more than 0.70 shown in bold, and VAF from 0.30 (SNVs) or 0.20 (INDELs) to 0.70 underlined.
